# Reducing Merkel cell activity in the whisker follicle disrupts cortical encoding of whisker movement amplitude and velocity

**DOI:** 10.1101/2022.07.08.499358

**Authors:** Clément E. Lemercier, Patrik Krieger

## Abstract

Merkel cells (MCs) and associated primary sensory afferents of the whisker follicle-sinus complex robustly code whisker self-movement, angle, and whisk phase during whisking. However, direct evidence of their roles in encoding whisker movement at cortical level is currently missing. To this end, spiking activity of primary somatosensory barrel cortex (wS1) neurons was measured in response to varying whisker deflection amplitude and velocity in transgenic mice with previously established reduced mechanoelectrical coupling at MC-associated afferents. Under reduced MC activity, wS1 neurons exhibited increased sensitivity to whisker deflection. This appeared to arise from a lack of variation in response magnitude to varying whisker deflection amplitude and velocity. This latter effect was further indicated by weaker variation in the temporal profile of the evoked spiking activity when whisker deflection amplitude and velocity varied. Nevertheless, under reduced MC activity, wS1 neurons retained the ability to discriminate stimulus features based on the timing of the first post-stimulus spike. Collectively, results from this study suggest that MCs contribute to both cortical encoding of whisker amplitude and velocity predominantly by tuning cortical response magnitude and by patterning evoked spiking activity, rather than in tuning cortical response latency.

## 1. Introduction

Rodents actively use their whiskers to explore their environment, discriminate objects and interact socially with conspecifics. The angle, speed and bending features of the whiskers, enable the inferring of information about the physical characteristics of objects being explored (Diamond et al., 2008). To detect and perceive a large variety of tactile features distinct classes of low threshold mechanosensory receptors (LTMRs) innervating the whisker follicle-sinus complex (FSC) (Ebara et al., 2002; Fundin et al., 1994; Rice et al., 1986, 1997; Tonomura et al., 2015) translate mechanical forces of whisker movement into trains of action potentials along primary trigeminal afferent neurons (Abraira and Ginty, 2013, Furuta et al., 2020; Gottschaldt et al., 1973; Severson et al., 2017). The neural code of whisker movement from neurons of the trigeminal ganglion (Jones et al., 2004; Szwed et al., 2003) travel next sequentially in a somatotopic manner through the brainstem trigeminal nuclei and somatosensory thalamic nuclei until the primary somatosensory barrel cortex (wS1) where information is integrated (Adibi et al., 2019; Bosman et al., 2011; Krieger and Groh 2015; Petersen, 2007; Sakurai et al., 2013). While knowledge about sensory processing along the trigemino-thalamo-cortical pathways is progressing, the precise functions played by the different whisker mechanoreceptor subtypes in cortical encoding whisker movement features are presently not fully understood.

Whisker mechanoreceptors have been classified accordingly to their morphological features into: Merkel, lanceolate, club-like, Ruffini endings (Ebara et al., 2002; Furuta et al., 2020; Rice et al., 1986, 1997; Tonomura et al., 2015) and as well as slowly or rapidly adapting encoders (SA and RA), according to their rates of adaptation to sustained mechanical stimulus (Gottschaldt et al., 1973). The combination of their morphological, positional, and discharge features are factors determining their properties in encoding diverse features of whisker movement (Furuta et al., 2020; Gottschaldt et al., 1973). Among the most prominent whisker mechanoreceptors is the Merkel cell-neurite complex (Gottschaldt and Vahle-Hinz, 1981; Halata et al., 2003; Iggo and Muir, 1969; Woo et al., 2015) in which mechanically excitable cells (i.e. Merkel cells, MCs) synaptically excite primary trigeminal afferent neurons to fire SA impulses (Chang et al., 2016; Higashikawa et al., 2019; Hoffman et al., 2018; Ikeda et al., 2014; 1994; Maksimovic et al., 2014; Maricich et al., 2009; Nakatani et al., 2015; Woo et al., 2014;2015). Activity of MCs tightly correlate with whisker displacement amplitude (Ikeda et al., 2014) and MC-associated afferents show robust coding of whisker self-movement, angle, and whisk-phase during whisking (Furuta et al., 2020; Severson et al., 2017). Considering that MCs constitute the primary sites for translation of whisker movement at SA afferents and that associated first-order trigeminal ganglion SA neurons encode whisker movement amplitude and velocity respectively by their response magnitude and latency (Kwegyir-Afful et al., 2008; Lottem et al., 2015; Shoykhet et al., 2000; Stüttgen et al., 2008), a reduction in MC activity is hypothesized to affect cortical encoding of these two modalities.

To assess this possibility, wS1 neurons were recorded juxtacellularly in anesthetized transgenic mice with established reduced mechanoelectrical coupling at MC-associated SA primary sensory afferents (Hoffman et al., 2018). Whiskers were deflected with six different deflection paradigms involving 3 different plateau amplitudes and 3 different ramp velocities and the coding properties of wS1 neurons were analysed and compared to littermate control mice. Data from this study suggest that MCs contribute to both cortical encoding of whisker deflection amplitude and velocity predominantly by tuning cortical response magnitude and by patterning evoked spiking activity, rather than tuning cortical response latency.

## 2. Materials and methods

### 2.1. Animals

Transgenic mice expressing tetanus neurotoxin light-chain subunit (TeNT) in MCs were obtained by crossing hemizygous *K14*^*Cre*^ male with homozygous *Rosa26*^*floxstopTeNT-GFP*^ female mice as described in Hoffman et al., (2018). Genotyping was performed at the Department of Physiology & Cellular Biophysics, Columbia University, New York, USA. Experiments involved 6 adult mice of both sexes including three littermate control (*Rosa26*^*floxstopTeNT-GFP*^) and three mice expressing TeNT in MCs (*K14*^*Cre*^*;R26*^*TeNT*^). Animal experimentation was conducted according to both European Union and German animal welfare regulations, and was approved by the local government ethics committee (Landesamt für Natur, Umwelt und Verbraucherschutz, Nordrhein-Westfalen, Germany).

### 2.2. Surgery and recording

Mice were anesthetized by an intraperitoneal injection of a saline mixture of urethane (1.2 - 1.4 g/kg, Sigma-Aldrich, Germany) and acepromazine (0.5 mg/kg). Body temperature was monitored and maintained at 37°C by the use of a closed loop heating pad (FHC, ME, USA). Oxygen was supplied continuously and the breathing rate was monitored on an oscilloscope connected to a piezoelectric disc (27 mm diameter) placed beneath the animal torso (Zehendner et al., 2013). When necessary, anesthesia was maintained with a supplementary injection of urethane (0.1 to 0.15 g/kg). The scalp was locally anesthetized with bupivacaine (0.25%) and incised. A head plate was fixed to the cranium with dental acrylic, the animal’s head was stabilized, and a craniotomy of ∼1mm2 was made over the wS1 cortex at 1.8 mm posterior to bregma and 3 mm lateral to the midline. The bath solution that was placed on top of the craniotomy comprised sterile physiological saline (0.9% NaCl). Juxtacellular recordings from wS1 neurons of head-fixed mice were made with 4-6 MΩ patch pipettes pulled from borosilicate filament glass (Hilgenberg GmbH, Germany; outer diameter: 1.5 mm, inner diameter: 0.86 mm) with a Sutter P-1000 puller (Sutter Instruments, Novato, CA, United States) filled with an extracellular solution containing 135 NaCl, 5.4 KCl, 1.8 CaCl_2_, 1 MgCl_2_, 5 HEPES (mM, pH ∼7.2). Signals were amplified (AxoClamp 2B amplifier, Axon Instruments), high-pass filtered at 300 Hz and sampled at 20 kHz (DigiData 1300, Axon Instruments) and visualized using pClamp 8 software (Molecular Devices, San Jose, CA, United States).

### 2.3. Whisker deflection paradigms

Whiskers were all trimmed to the same length of ∼1 cm to ensure equal movement when deflected. Once a neuron was approached, a thin wooden stick (diameter: 2 mm) was used as a probe to manually and individually deflect the whiskers, and an audio monitor was used to identify the whisker that evokes the strongest response with shortest latency (i.e. the principal whisker, PW). The PWs were deflected using a ramp-and-hold movement by inserting their tips into a glass capillary (placed ∼1 mm from the muzzle) glued to a piezo wafer (PL127.10; PI Ceramics, Lederhose, Germany) controlled by a piezo filter and amplifier (Sigmann Elektronik, Hüffenhardt, Germany). The plateau amplitude (i.e. displacement) of the piezo wafer was monitored with a video camera (USB8MP02G-SFV, Shenzhen Ailipu Technology Co., Ltd) and peak ramp velocity was estimated as the ratio between piezo plateau amplitude to piezo-amplifier transition time. The maximal peak ramp velocity was thus estimated to be ∼0.6 mm/ms to the maximal plateau amplitude achievable of ∼1.2mm, which considers displacement due to ringing. Spiking activity of wS1 neurons was measured across six distinct whisker deflection paradigms, including three paradigms with varying plateau amplitude at constant ramp velocity (displacement of ∼1.2, ∼0.5 and ∼0.25 mm at a velocity of ∼0.6 mm/ms) and three paradigms with varying ramp velocity at constant plateau amplitude (velocity of ∼0.6, ∼0.12 and ∼0.06 mm/ms at a displacement of ∼1.2 mm). The holding time of the piezo wafer was set at 100 ms. Deflection paradigms were presented as a random sequence of recording blocks comprising 20 stimuli delivered at 0.5 Hz. For each deflection paradigms, spiking activity was monitored over 3 recording blocks (i.e. 60 deflections per deflection paradigms). An additional paradigm was designed to evaluate change in the response adaptation to repetitive whisker deflection and comprised trains of 8 deflections at 8 Hz (displacement of ∼1.2 mm at a velocity of ∼0.6 mm/ms, piezo wafer holding time set at 25 ms) delivered at 0.2 Hz over 3 recording blocks of 20 sweeps.

### 2.4. Histology

At the end of the experiment, animals were overdosed with urethane (3 g/kg), transcardially perfused with paraformaldehyde 4% (V/V in phosphate buffer) and their brains were removed and prepared for flattened corticotangential sectioning, as described by Lauer et al., (2018). Cortical sections were stained for cytochrome oxidase, and layer IV barrel-field cytoarchitecture was visualized under bright-field light microscopy. Mystacial pads were dissected, cryoprotected in 0.1M phosphate buffer solution containing 30% Sucrose (w/v) and cut into 50 μm thick transversal sections with a cryostat microtome (CM3050S, Leica Biosystems). Sections were counterstained with the nuclear dye 4′,6-Diamidine-2′-phenylindole dihydrochloride (DAPI, Sigma-Aldrich, Germany) and actin filaments were labeled with ATTO 647-phalloidin conjugate (Hypermol EK, Germany). Optical sections of the whisker follicle-sinus complex were acquired by confocal-like optical sectioning with an Apotome Microscope (Axioimager Z1, ZEISS Microscopy, Germany). The expression of TeNT-GFP in *Cre-*positive mice was confirmed by the presence of a green fluorescence signal in the MC-dense region of the whisker FSC.

### 2.5. Data analysis

Spike detection was done with the template search function of the software pClamp 11.2 (Molecular Devices, San Jose, CA, United States), and spontaneous spiking and peri-stimulus spike time-stamps were determined. For each recorded neuron, spontaneous spiking activity was estimated over segments of 1 sec distantly located from deflection onset (1 sec). To assess the overall effect of reducing MC activity on evoked cortical spiking activity, the number of evoked spikes, response probability (whether spikes were elicited or not) and first-spike latency to whisker protraction were firstly measured on a 100 ms response window. Response adaptation to repetitive whisker deflection (including responses to protraction and retraction) was evaluated by quantifying and comparing the number of evoked spikes at each deflection on a 100 ms response window. Changes in response magnitude (i.e. evoked spike count) to varying whisker protraction amplitude or velocity were quantified on a shorter response window of 30 ms. Peristimulus time histograms (PSTH) were determined over a window of 150 ms (10 ms bin size) including 50 ms prior to and 100 ms after stimulus onset. Cumulative distribution functions (CDFs) derived from PSTHs were used to analyze the temporal aggregation of the evoked spiking activity. Differences between CDFs in response to varying whisker deflection intensity (either amplitude or velocity) were evaluated by plotting CDFs at intermediary (*y*) or minimal (*y*’) deflection intensity against CDFs at maximal (*x*) deflection intensity (i.e. *y = f(x)*) and differences were quantified in a pooled manner (across *y* and *y*’) with the following equation:

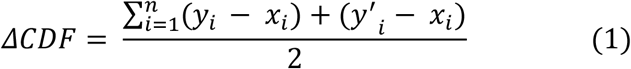

### 2.6. Statistics

Statistical analysis was performed using the software R (R Core Team 2021) and the ‘PMCMR-plus’ library (Pohlert 2020). To report statistics in text, control and mice expressing TeNT in MCs are respectively abbreviated as ‘CON’ and ‘MC_TeNT_’. Differences between and within experimental groups were respectively evaluated with a Wilcoxon rank sum test and with a Friedman test. A *p*-value < 0.05 was considered statistically significant. Unless specified, data are reported in the text as Median and Interquartile range (IQR, .i.e. the differences between Q3 and Q1). Sample size (*n*) is reported in the figure legends. Figures were produced with the R package ‘ggplot2’ (Wickham, 2016).

## 3. Results

### 3.1. Expression of TeNT in the MC-dense region of the whisker FSC and barrel-field cytoarchitecture

To reduce mechanoelectrical coupling at Merkel endings, the vesicular release machinery of MCs was blocked by expressing TeNT in cells expressing K14 epidermal precursor (Hoffman et al., 2018; Morrison et al., 2009; Van Keymeulen et al., 2009; Woo et al., 2010; Yamamoto et al., 2003). Genetic expression of TeNT-GFP in the whisker FSC was confirmed by green fluorescence signal located in the MC-dense region (MDR) at the level of the ring sinus (RS) between the ringwulst (RW) and the inner canonical body (ICB) (Ebara et al., 2002; Rice et al., 1986; Whiteley et al., 2015) (Fig 1A-1B). The effect of reducing MC activity on layer IV wS1 cytoarchitecture was evaluated on flattened corticotangential sections stained for cytochrome oxidase (Land and Simons, 1985; Lauer et al., 2018; Welker and Woolsey, 1974) (Fig 1C-1D). A reduction in MC activity was not associated to a noticeable change in individual barrels size from respectively identified barrel rows (observation from one slice of both group).

**Fig 1.**
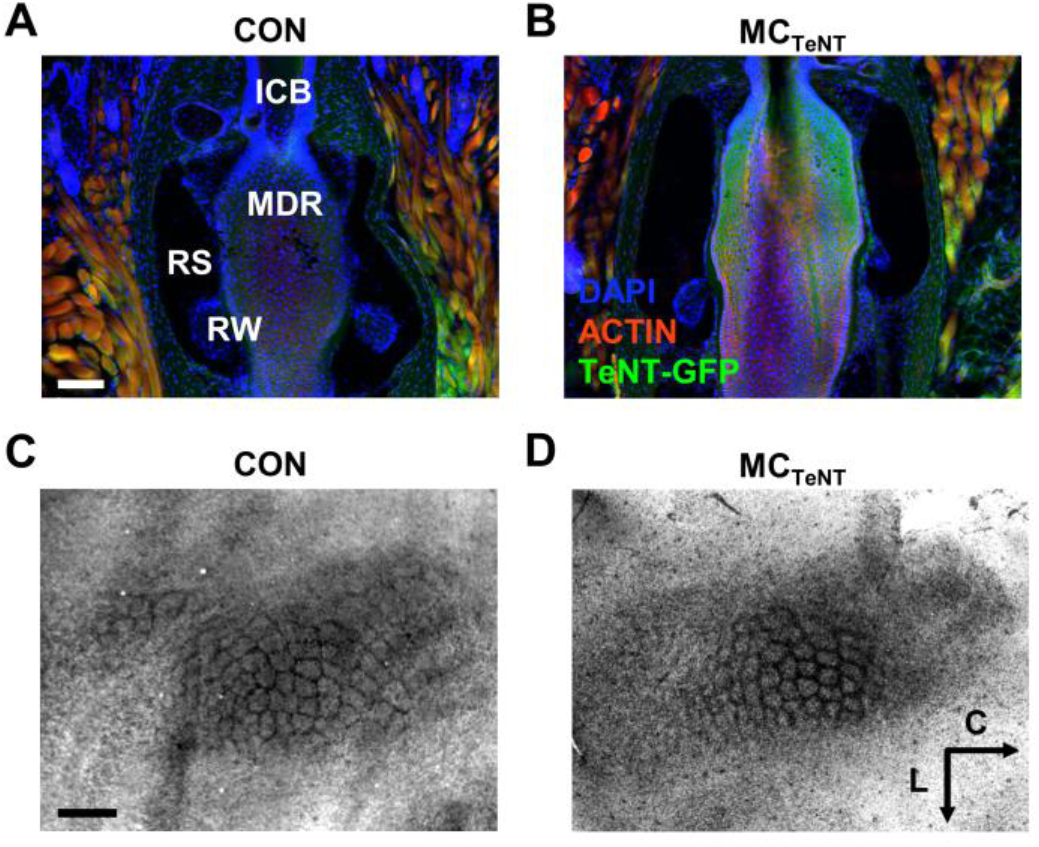
Expression of TeNT in the MC-dense region of the whisker FSC and barrel-field cytoarchitecture. **(A-B)** Longitudinal sections of whisker FSC stained for cell nuclei and actin filaments from control (A) and MC_TeNT_ (B) mice. Note the presence of a green fluorescence signal indicative of the expression of TeNT-GFP at the level of the MC-dense region (MDR) of the whisker FSC in MC_TeNT_ mice. ICB: Inner canonical body; MDR: Merkel cell dense region; RS: Ring sinus; RW: Ringwulst. Scale bar = 100 μm. **(C-D)** Flattened tangential sections of layer IV wS1 cortex stained for cytochrome oxidase from control (C) and MC_TeNT_ mice (D). Scale bar = 500μm, L: lateral, C: caudal.

### 3.2. Reducing MC activity increases wS1 sensitivity to whisker deflection

The median depth, relative to the pia mater, of the recorded wS1 neurons was similar between the two experimental groups (CON = 494 μm (230), n = 9; MC_TeNT_ = 605 μm (177), n = 10; *p* = 0.4967 - Fig 2A). The effect of reducing MC activity on whisker-evoked wS1 spiking activity was first evaluated by pooling the data collected across the six different whisker deflection paradigms with varying amplitude or velocity.

**Fig 2.**
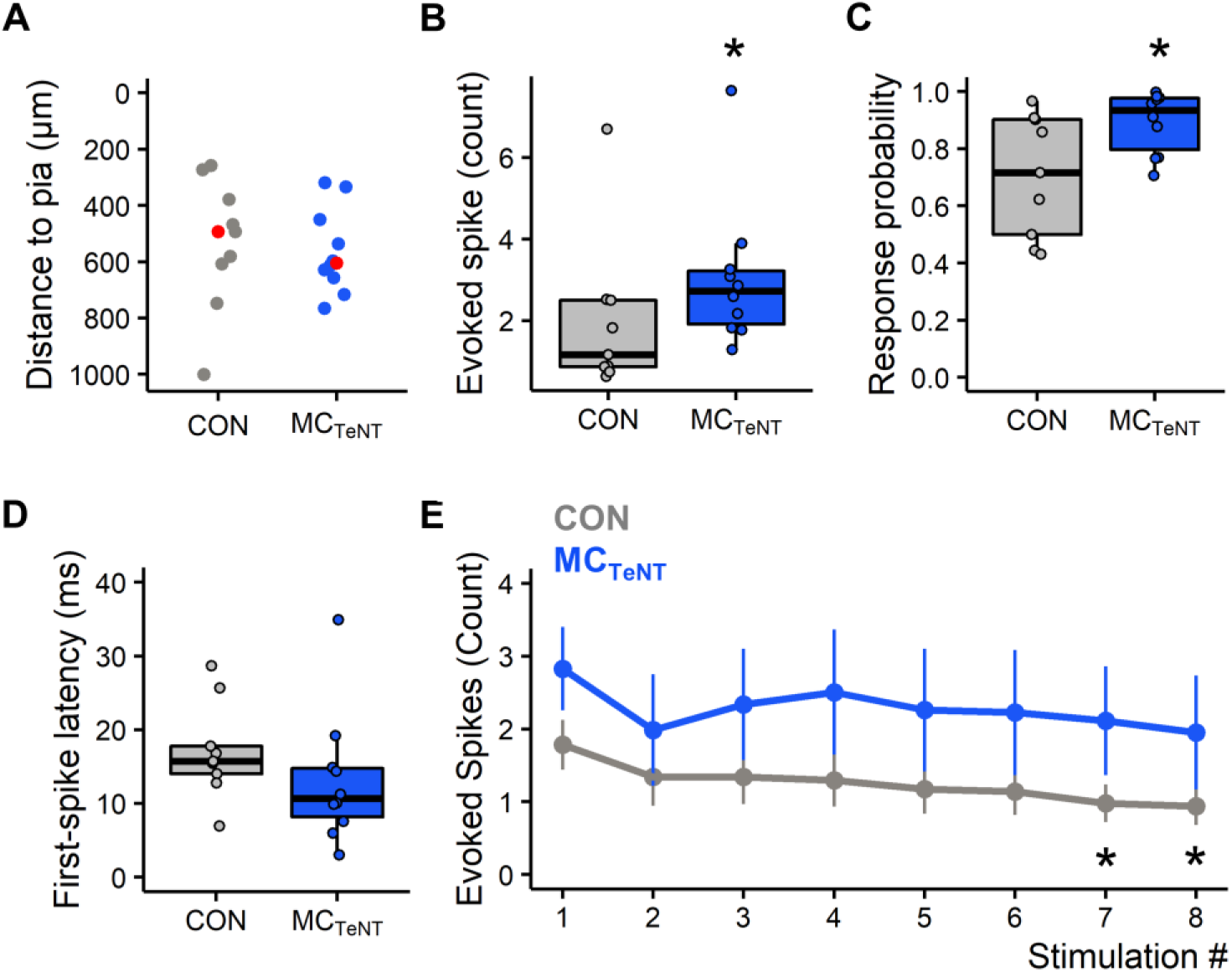
Overall effect of reducing MC activity on whisker-evoked wS1 spiking, response probability, first-spike latency, and response adaptation to repetitive deflections. **(A)** Relative depth distribution to pia of the recorded wS1 neurons (CON: n = 9; MC_TeNT_: n = 10), red markers indicate the median recording depth. **(B-D)** Boxplots showing the grand-average effect (collapsed across the six different whisker stimulation paradigms) of reducing MC activity on the number of whisker-evoked spikes, response probability and first-spike latency (over 100 ms response window) (CON: n = 9; MC_TeNT_: n = 10). **(E)** Average number of evoked spikes along 8 repetitive whisker deflections delivered at 8 Hz (CON: n = 7; MC_TeNT_: n = 7), data shown as Mean ± SEM. (*) Indicate significant differences in control between 1^st^/7^th^ and 1^st^/8^th^ deflection pairs. (A-D) Wilcoxon rank sum test. (E) Friedman test followed by Conover’s all-pairs posthoc test.

Although expression of TeNT in MCs was previously reported to reduce mechanoelectrical coupling at SA afferents by about ∼50 % (Hoffman et al., 2018) our data indicate that reducing MC activity increased wS1 neurons sensitivity to whisker deflection (Fig 2B-2C). This effect manifested in an increase in the number of whisker-evoked spikes (CON = 1.17 spikes (1.63); MC_TeNT_ = 2.72 spikes (1.30); *p* = 0.0349 - Fig 2B) and in response probability (CON = 0.72 (0.40); MC_TeNT_ = 0.93 (0.18); *p* = 0.0279 - Fig 2C) but occurred without significant change in first-spike latency (CON = 15.74 ms (3.73); MC_TeNT_ = 10.66 ms (6.61); *p* = 0.1333 - Fig 2D) and in spontaneous spiking activity (CON = 2.17 Hz (5.14); MC_TeNT_ = 4.65 (4.83); *p* = 0.2775 - data not shown).

The increased sensitivity to whisker deflection was accompanied by a lack of response adaptation to repetitive whisker deflections. Indeed, response adaption was significantly observed in control between the 1^st^/7^th^, and 1^st^/8^th^ deflection pairs (1^st^ = 1.78 spikes (0.34), 7^th^= 0.98 (0.26), 8^th^ = 0.94 (0.26), data expressed as Mean (SEM); 1^st^/7^th^: *p* = 0.0300; 1^st^/8^th^: *p* = 0.0214 - Fig 2E) but not under conditions of reduced MC activity (1^st^ = 2.83 spikes (0.57), 7^th^ = 2.11 (0.75), 8^th^ = 1.95 (0.78), data expressed as Mean (SEM); 1^st^/7^th^: *p* = 0.3574; 1^st^/8^th^: *p* = 0.6384 - Fig 2E), although spiking activity tended on average to decrease.

### 3.3. Reducing MC activity alters the response magnitude of wS1 neurons to varying whisker deflection amplitude and velocity

wS1 neurons exhibit responses magnitude and timing that generally depend on whisker deflection amplitude and velocity (Ito and Kato, 2002; Pinto et al., 2000; Wilent and Contreras, 2004). Decreasing whisker deflection amplitude or velocity decreased the evoked spiking activity in control but not in condition of reduced MC activity (varying amplitude: CON, *p* = 0.0030; MC_TeNT_, *p* = 0.1496; varying velocity: CON, *p* = 0.0207; MC_TeNT_, *p* = 0.2765) – (Fig 3A-3B and S1 Table). Under reduced MC activity, the coding properties of wS1 were partially preserved, as seen in the decreased response probability when deflection amplitude was reduced (CON, *p* = 0.0017; MC_TeNT_, *p* = 0.0122) – (S1 Table) and with increased first-spike latency when deflection amplitude or velocity was reduced (varying amplitude: CON, *p* = 0.0043; MC_TeNT_, *p* = 0.002; varying velocity: CON, *p* < 0.001; MC_TeNT_, *p* = 0.0025) – (S1 Table). Nevertheless, response probability and first-spike latency at low deflection amplitude or low velocity were on average higher and faster than in control (S1 Table). These data suggest that MCs contribute to both cortical encoding of whisker movement amplitude and velocity predominantly by tuning wS1 response magnitude than firing latency and probability.

**Fig 3.**
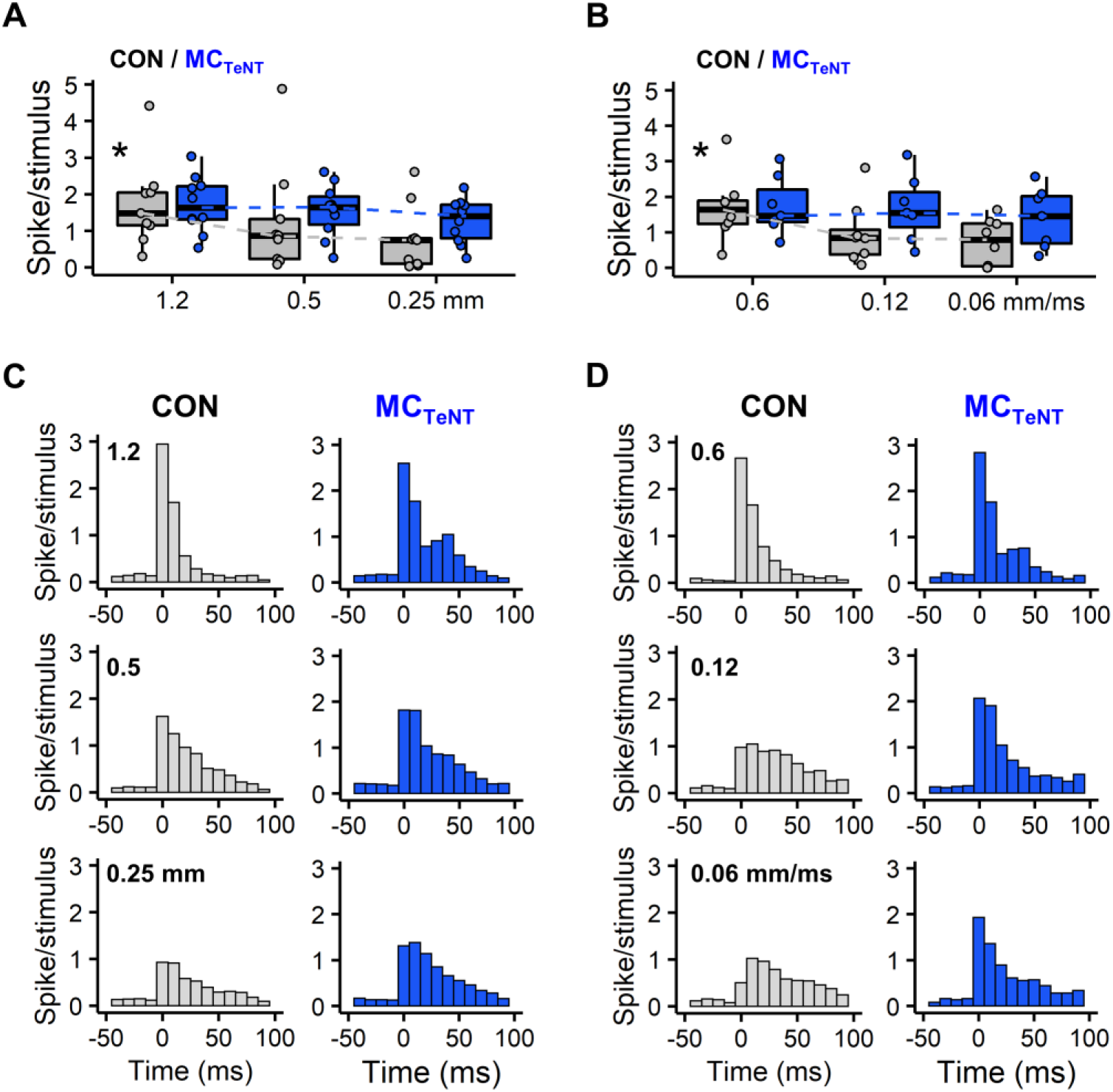
Effect of reducing MC activity on whisker-evoked wS1 spiking activity in response to varying whisker deflection amplitude or velocity. (**A-B)** Average number of evoked spikes (over 30 ms response window) across the different whisker deflection paradigms with varying stimulation amplitude (A) or velocity (B). Data from MC_TeNT_ mice are displayed in blue. (*) Indicates that spiking activity in control significantly decrease when deflection amplitude or velocity varied (Friedman test). Dashed lines link medians. **(C-D)** Population PSTHs (average response over 60 sweeps, bin width 10 ms) for each deflection paradigms with varying amplitude (C) or velocity (D). (A-D) Varying amplitude: CON, n = 9; MC_TeNT_, n = 10 - Varying velocity: CON, n = 8; MC_TeNT_, n = 7.

In controls, inspection of PSTHs indicates that variations in whisker deflection amplitude or velocity influence patterns of evoked spiking activity. Effects varied from patterns with rather fixed timing over a short period at strong deflection intensities, through patterns with more variable timing over a longer period at weaker deflection intensities (Fig 3C-3D). Interestingly, PSTHs under reduced MC activity, especially for varying deflection velocity, appear to exhibit less change in the temporal aggregation of evoked spiking activity than in controls (Fig 3C-3D). To evaluate changes in whisker-evoked spike patterns, cumulative distribution functions (CDFs) were derived from PSTHs, and changes in CDFs at intermediary and minimal deflection intensity (either amplitude or velocity) were assessed against CDFs at maximal deflection intensity used as reference (Fig 4, see materials and methods).

**Fig 4.**
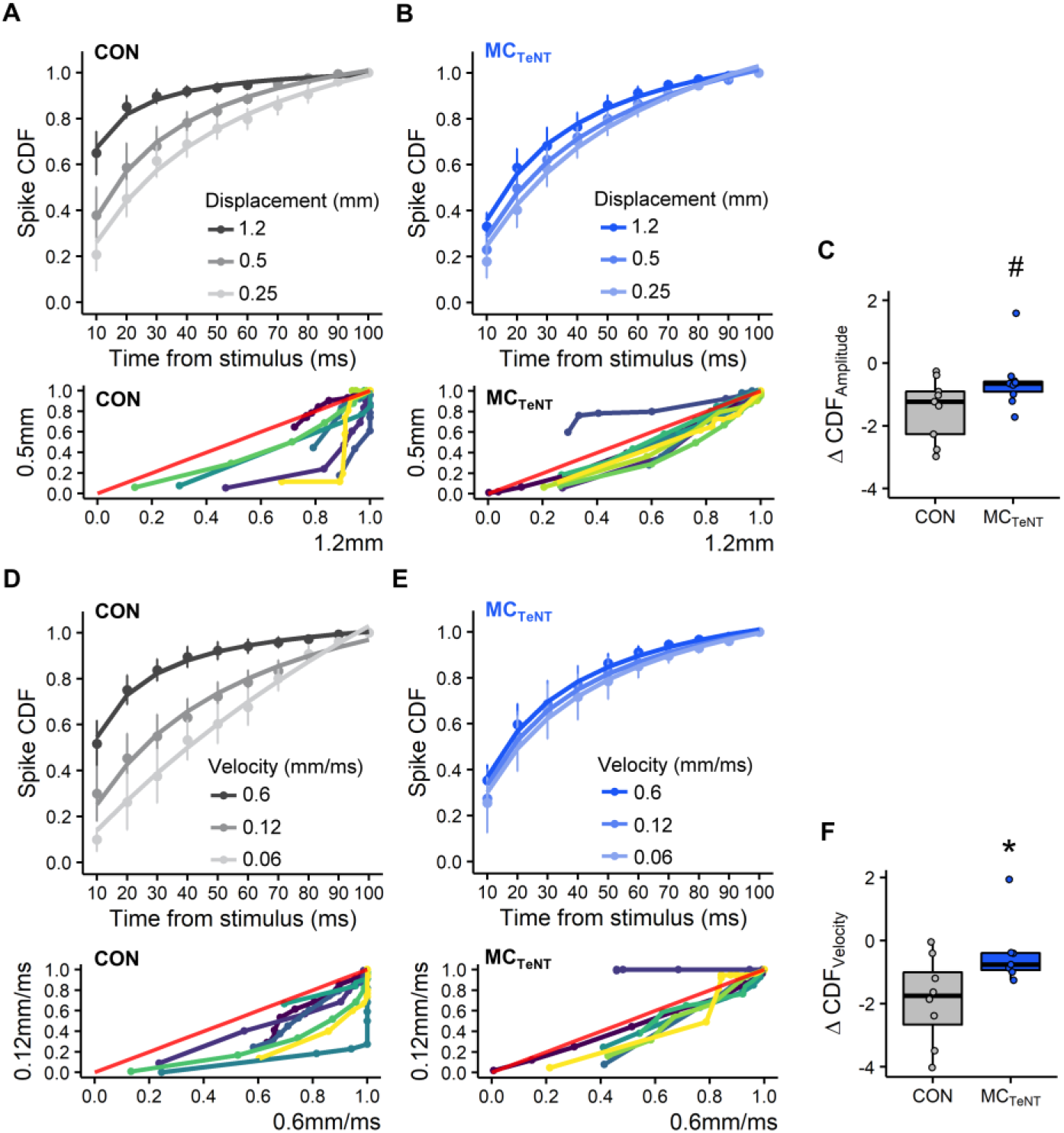
Effect of reducing MC activity on the temporal profile of evoked spiking activity in response to varying whisker deflection amplitude or velocity. **(A-B)** Change in spike CDFs in response to varying whisker deflection amplitude from control (A) and MC_TeNT_ mice (B). Below is shown the crossed-CDF variation between maximal and intermediary deflection amplitude for each individual neurons; red line denotes an identical distribution (i.e. *f(y) = f(x)*). **(C)** Pooled variation in CDFs across maximal/intermediary and maximal/minimal whisker deflection amplitude, # is indicative of *p* = 0.0563. **(D-F)** Identical panel structure to (A-C) for data obtained with varying whisker deflection velocity. (A-B and D-E) data shown as Mean ± SEM. Fits computed with non-linear least squares regression, Pearson’s Goodness-of-Fit: *p* < 0.05. (C and F) Wilcoxon rank sum test. (A-F) Varying amplitude: CON, n = 9; MC_TeNT_, n = 10 - Varying velocity: CON, n = 8; MC_TeNT_, n = 7.

In controls, CDF rates of evoked spiking activity decreased, as deflection amplitude or velocity decreased (Fig 4A and 4D), indicating that lowering whisker deflection intensity causes spiking activity to aggregates less suddenly and more distributedly in time. Shifts in CDF rates across whisker deflection intensities were on average weaker in condition of reduced MC activity (Fig 4B and 4E) indicating that spiking activity tended to aggregates equally in time, independently of changes in whisker deflection amplitude or velocity. In respect to control, differences in CDFs for varying whisker deflection amplitude or velocity were weaker under reduced MC activity (varying amplitude: *p* = 0.0563 - Fig 4C; varying velocity: *p* < 0.0360 - Fig 4F). Collectively these data suggest that beyond contributing to wS1 response magnitude, MCs also contribute to the patterning of wS1 spiking activity.

## 4. Discussion

Data from this study suggest that reducing MC activity increases wS1 neurons sensitivity and response probability to whisker deflection. As activity of MCs and associated afferents tightly correlate with whisker movement during passive and active touch (Furuta et al., 2020; Hoffman et al., 2018; Ikeda et al., 2014; Severson et al., 2017), a reduction in their activity was presumed to produce the opposite effect on wS1 activity. Nevertheless, increased cortical sensitivity to cutaneous touch of glabrous skin has been recently reported in the somatosensory cortex (S1) in response to genetic ablation of MCs (Emanuel et al., 2021). The interpretation for the shift in S1 sensitivity was that MC-associated afferents recruit subcortical elements involved in setting S1 sensitivity. Another interpretation, albeit complementary but more speculative, comprises a role of MC-associated afferents in gating early sensory inputs from other LTMRs by interacting potentially with the inhibitory circuitry of the trigeminal ganglion (Hayasaki et al. 2006).

In the sample of recorded neurons, increased sensitivity to whisker deflection was suggested to occur with a lack of response adaptation to repetitive deflection. Although spiking activity to repetitive deflection decreased, on average, in conditions of reduced MC activity, this effect did not reach significance. Low sample size and higher spiking variability under conditions of reduced MC activity are most likely to be the cause. While neural adaptation along the whisker-barrel neuraxis occurs at all stages of sensory processing (Adibi and Lampl, 2021), primary sensory neurons of the trigeminal ganglion exhibit little adaptation to repeated whisker deflection in contrast to subsequent sensory relays (Adibi and Lampl, 2021; Ganmor et al., 2010). Adaptation appears to be rather inherited in subsequent sensory relays, making the absence of sensory adaptation taking place at the level of the cortex difficult to interpret under conditions of reduced MC activity. Additional experiments will allow to better test this observation.

The data of the present study suggest that MCs contribute to cortical encoding of both whisker deflection amplitude and velocity predominantly by tuning cortical response magnitude and by patterning the evoked spiking activity. Nevertheless, under conditions of reduced MC activity, cortical encoding of whisker deflection amplitude and velocity was still preserved, notably in terms of changes in first-spike latency and response probability. Cortical first-spike latency thus may rely on other RA whisker LTMRs subtypes, such as lanceolate- and club-like endings. Indeed, these two LTMRs subtypes exhibit faster response latency than SA Merkel-endings (Furuta et al., 2020). Nevertheless, it has been recently reported that depending on their location in the whisker FSC (i.e. deep vs. superficial), Merkel-endings exhibit different adapting properties (i.e. SA vs. RA) and therefore differential sensitivities to whisker movement (Furuta et al., 2020). Therefore, in the present study, the reported cortical effects are assumed to arise from a reduction in mechanoelectrical coupling at both types of Merkel endings.

Whisker movement amplitude and velocity are features used by rodents for inferring information about their environment (Arabzadeh et al., 2005; Pammer et al., 2013). Along the whisker pathway, whisker movement amplitude and velocity are respectively better represented by response magnitude and timing (Bale et al., 2015; Kwegyir-Afful et al., 2008; Lottem et al., 2015; Shoykhet et al., 2000) although wS1 neurons are more sensitive to whisker movement velocity than amplitude (Pinto et al., 2000). Analysis of coding strategies of whisker movement at the level of the cortex indicated that the majority of the information about the stimulus identity is retained in the timing of the first post-stimulus spike (Bale and Peterson, 2009; Panzeri et al., 2001; Petersen et al., 2001). Therefore, as previously reported with genetic ablation of MCs (Maricich et al., 2012), our data suggest that mice with reduced MC activity can still rely on their whiskers through other LTMRs subtypes to perform tactile behaviors such as texture discrimination tasks. In addition of being the main organ for tactile perception in mouse, whiskers self-motion combined to those of facial hairs was proposed to contribute to mouse facial proprioception (Severson et al., 2017, 2019). Considering the ability of MC-associated afferents to signal both static and whisker self-movement in their rates of sustained discharge, this LTMR might constitutes a good candidate in contributing to mouse facial proprioception.

## 5. Conclusion

The major outcome of this preliminary study indicates that MCs in the whisker FSC contribute to both cortical encoding of whisker movements amplitude and velocity predominantly by tuning wS1 response magnitude and by patterning the evoked spiking activity. As wS1 neurons retained the ability to discriminate stimulus features based on the timing of the first post-stimulus spike, MCs are thus suggested to play secondary roles over other LTMRs in tuning cortical response latency. While the present study was able to replicate a recent finding on the role played by MCs in tuning S1 sensitivity in response to cutaneous touch (Emanuel et al., 2021), its results and interpretation were based on a relatively sample size. Further investigations are thus required to confirm the proposed roles played by MCs in cortical encoding of whisker movement amplitude and velocity.

## Abbreviations

CDF: cumulative distribution function
FSC: follicle-sinus complex
IQR: interquartile range
K14: keratin 14
LTMR: low threshold mechanoreceptor
MC: Merkel cell
PSTH: peristimulus time histograms
PW: principal whisker
RA: rapidly adapting
SA: slow adapting
TeNT: tetanus neurotoxin light-chain subunit
wS1: primary somatosensory barrel cortex.

## Acknowledgments

The authors thank Prof. Ellen Lumpkin (Department of Physiology & Cellular Biophysics, Columbia University, New York, NY 10032) for the gift of the transgenic mouse lines and Ute Neubacher (Manahan-Vaughan lab, Department of Neurophysiology, Medical Faculty, Ruhr University Bochum) for helpful assistance in histology and imaging.

## Author contributions statement

C.E.L. and P.K. designed the research. C.E.L. performed the experiments, analyzed the data, and wrote the paper.

## Data accessibility statement

The data supporting this study can be made available upon reasonable request to the corresponding author.

## Supplementary material

**S1 Table.**
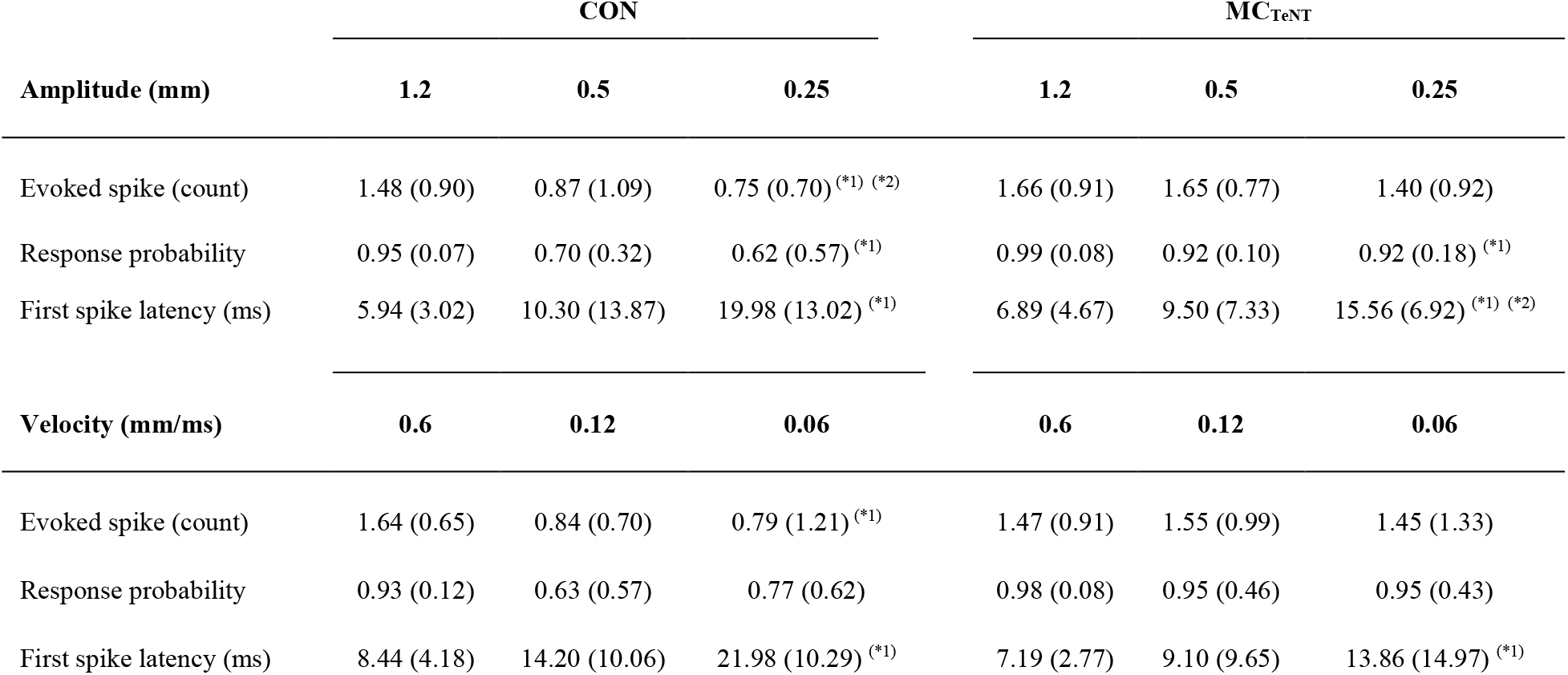
Effect of reducing MC activity on whisker-evoked wS1 spiking activity, response probability and first-spike latency in response to varying whisker deflection amplitude and velocity. Control and mice expressing TeNT in MCs are respectively abbreviated as ‘CON’ and ‘MCTeNT’. Data expressed as Median (IQR). Evoked spike count was determined over 30 ms response window. Amplitude: CON, n = 9; MCTeNT, n = 10; Velocity: CON, n = 8; MCTeNT, n = 7. (*1) Indicate significant difference (p < 0.05) between maximal and minimal deflection amplitude (1.2 / 0.25 mm) or velocity (0.6 / 0.06 mm/ms). (*2) Indicate significant difference (p < 0.05) between maximal and intermediary deflection amplitude (1.2 / 0.5 mm) or velocity (0.6 / 0.12 mm/ms). Friedman test followed by Conover’s all-pairs posthoc test.

